# Too hot to handle? The impact of the 2023 marine heatwave on Florida Keys coral

**DOI:** 10.1101/2024.08.31.610635

**Authors:** Karen L. Neely, Robert J. Nowicki, Michelle A. Dobler, Arelys A. Chaparro, Samantha M. Miller, Kathryn A. Toth

## Abstract

The marine heatwave in the summer of 2023 was the most severe on record for Florida’s Coral Reef, with unprecedented water temperatures and cumulative thermal stress precipitating near 100% coral bleaching levels. An existing SCTLD coral fate-tracking program assessed over 4200 coral colonies across five offshore and four inshore reef sites approximately every two months, allowing for analyses of bleaching-related mortality and diseases during and after the marine heatwave. Across the vast majority of assessed corals, including multiple sites and species, there was no partial or full mortality as a result of the 2023 bleaching event. The two sites that did experience substantial bleaching-related mortality were those experiencing the highest levels of cumulative thermal stress. However, the substantial acute mortality at one of them occurred at relatively low levels of cumulative stress, suggesting death was the result of exceeding thermal maxima. At the two sites with notable mortality, 43% and 24% of all monitored corals died, but mortality varied among species. Brain corals fared worse than boulder corals, with *Pseudodiploria strigosa* the most heavily impacted species. The health status of corals before the bleaching event had little impact on whether they experienced disease or bleaching-related mortality during the event. At three sites, we observed unusual lesions on *Orbicella faveolata* colonies shortly after color returned to the corals; the lesions were only observed for a few months but on some colonies led to substantial tissue loss. Though not part of the coral monitoring program, we also observed substantial losses and local extinctions of Acroporid corals at most sites, as well as probable local extinctions of octocorals at three of the four inshore reefs. Though most reef-building corals came through the 2023 event with no mortality, continually rising temperatures are likely to make these temperature regimes more common and widespread. We encourage future research on what the unusual *O. faveolata* lesions are, and why the brain and boulder corals fared differently at highly-impacted sites. Our results also provide perspective on how restoration strategies, particularly those focused on species likely to die under current and future climate regimes, should consider shifting focus to species likely to survive. Finally, these results highlight the importance of this type of monitoring, with a focus on fate-tracking individuals through disturbance events, including a large number of individuals of multiple species across a geographic range and multiple habitats.

## Introduction

Stony corals on tropical reefs are subject to a variety of stressors, and as a result of these, the majority of reefs worldwide are in some state of decline. The most geographically extensive of these stressors is coral bleaching, primarily resulting from increasingly warmer seas periodically exceeding corals’ preferred temperature thresholds. During these marine heatwaves (MHWs), the symbiosis between tropical scleractinian corals and their associated zooxanthellae symbionts breaks down (Brown, 1997), leaving the corals’ tissues transparent and the coral appearing white or “bleached.” Though the coral is still alive when bleached, without its symbionts it is deprived of much of its nutritional uptake (Muscatine et al., 1981), which ultimately results in death unless the stress event diminishes and the algal symbionts recolonize the coral. Bleaching events have become increasingly common across all tropical oceans (Glynn, 1993; Hoegh-Guldberg, 1999), and predictions for future frequency and intensity of the events are increasingly dire (Hoegh-Guldberg, 1999; McWilliams et al., 2005; Mellin et al., 2024).

Florida’s Coral Reef represents the northernmost extent of reef development in the Caribbean. Through 2023, eight region-wide mass-bleaching events have been documented across Florida Keys reefs. The 1997-98 bleaching events corresponded with reduction in species richness and coral cover (Somerfield et al., 2008). The back-to-back bleaching events in 2014 and 2015 led to some partial or total coral mortality at certain sites (Gintert et al., 2018) as well as losses to some stands of *Acropora palmata* (Neely et al., 2022) and *Dendrogyra cylindrus* (Neely et al., 2021a). However, other events and observations on other species showed minor or no losses; for example Fitt et al. (1993) found no mortality in their studied colonies from the 1987 event.

The intensity of past thermal stress events in Florida paled in comparison to the summer of 2023, which saw an unprecedented heat wave in the Florida Keys. On land, heat records were broken on 31 of the 62 days in July and August since record keeping began in 1950 (NOAA National Weather Service, 2023). NOAA’s Coral Reef Watch monitors sea surface temperatures, calculates cumulative thermal stress, and issues bleaching watches and warnings for reefs worldwide (Figure 1). A warning indicates that sea surface temperatures are above the local bleaching threshold, and that cumulative thermal stress is above 0 degree heating weeks (DHW) but less than 4 DHW. Bleaching alerts are classed into alert level 1, in which DHW are greater than 4 but less than 8, and alert level 2 in which DHW are greater than 8. In 2023, a bleaching warning was issued on June 14, an alert level 1 on July 6, and an alert level 2 on July 16. The alerts remained active through September 30 (NOAA Coral Reef Watch, 2023). This resulted in mass bleaching of all or nearly all scleractinian corals throughout the region, and some anecdotal and media reports suggested that this might lead to widespread coral mortality.

**Figure 1.**
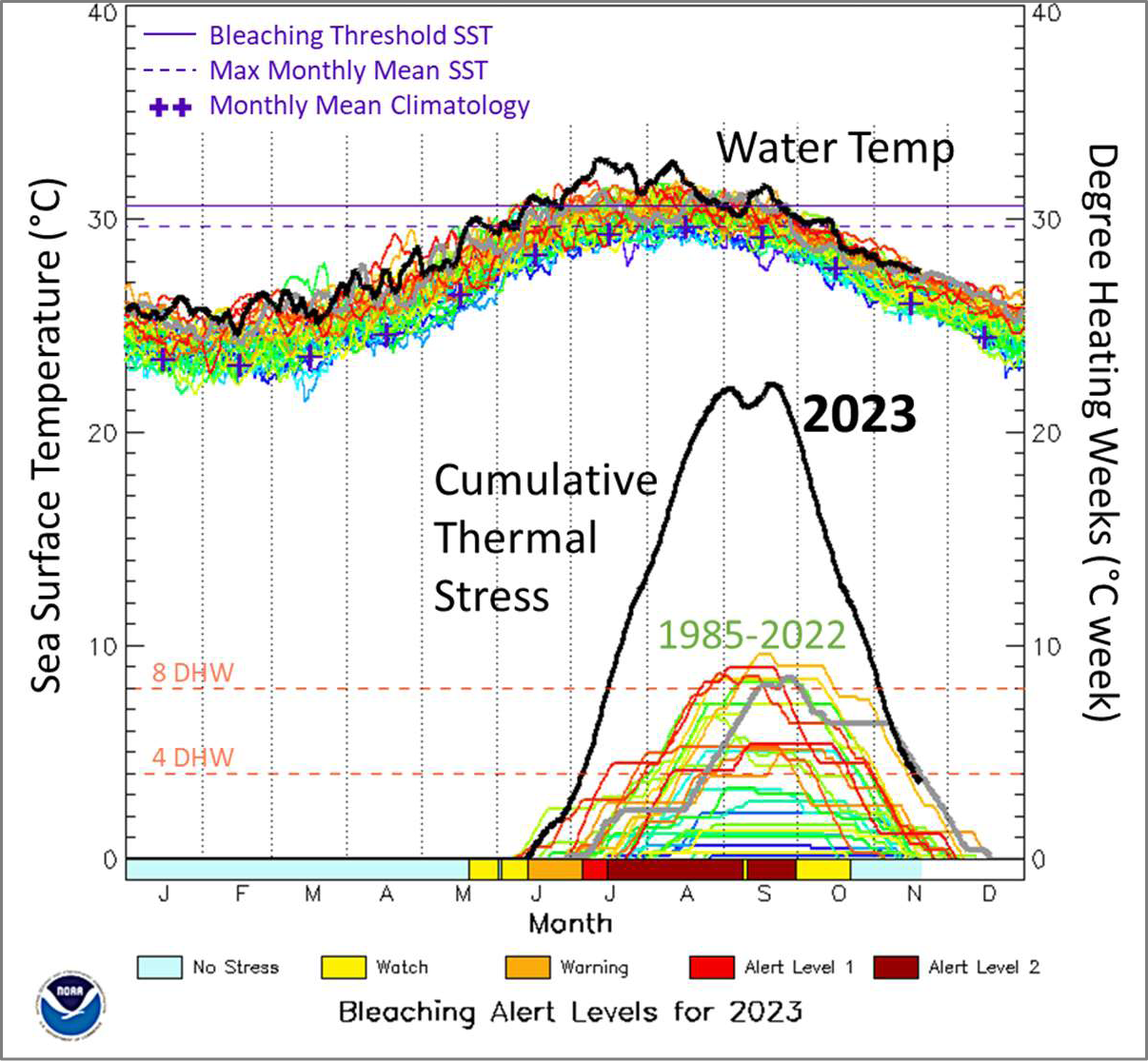
Temperature (top) and cumulative thermal stress degree heating weeks (bottom) data for the Florida Keys collected by NOAA Coral Reef Watch since 1985. Black lines show the values through the 2023 hyperthermal event; colored lines show each year’s data from 1985-2022. Colored bars along the bottom indicate the bleaching alert levels through 2023. Figure slightly modified from NOAA Coral Reef Watch.

Using an existing monitoring program on SCTLD-affected and treated corals throughout the Florida Keys, we quantified the impacts of this MHW on boulder and brain corals across inshore and offshore reefs throughout the Florida Keys.

## Methods

### Field assessments

From 2019-2024, stony corals at reef sites throughout the Florida Keys were monitored approximately every other month through a stony coral tissue loss disease (SCTLD) intervention program (Neely et al., 2021b). A total of five offshore and four inshore/midchannel patch reef sites were assessed before, during, and after the 2023 summer (Figure 2). Part of one reef (Cheeca Rocks) was also monitored an additional time during the peak bleaching impact to further assess temporal mortality patterns. Offshore sites were, from southwest to northeast, Sand Key, Looe Key, Sombrero Reef, Grecian/Key Largo Dry Rocks, and Carysfort. Inshore sites were Newfound Harbor, Marker 48, Cheeca Rocks, and Hen and Chickens. At offshore sites, the bases of corals were between 1.2 – 9.4 m in depth; at inshore sites, corals were between 1.2 – 6.4 m in depth. The number of corals regularly monitored at each site varied from 96 to 1335 (Table 1).

**Figure 2.**
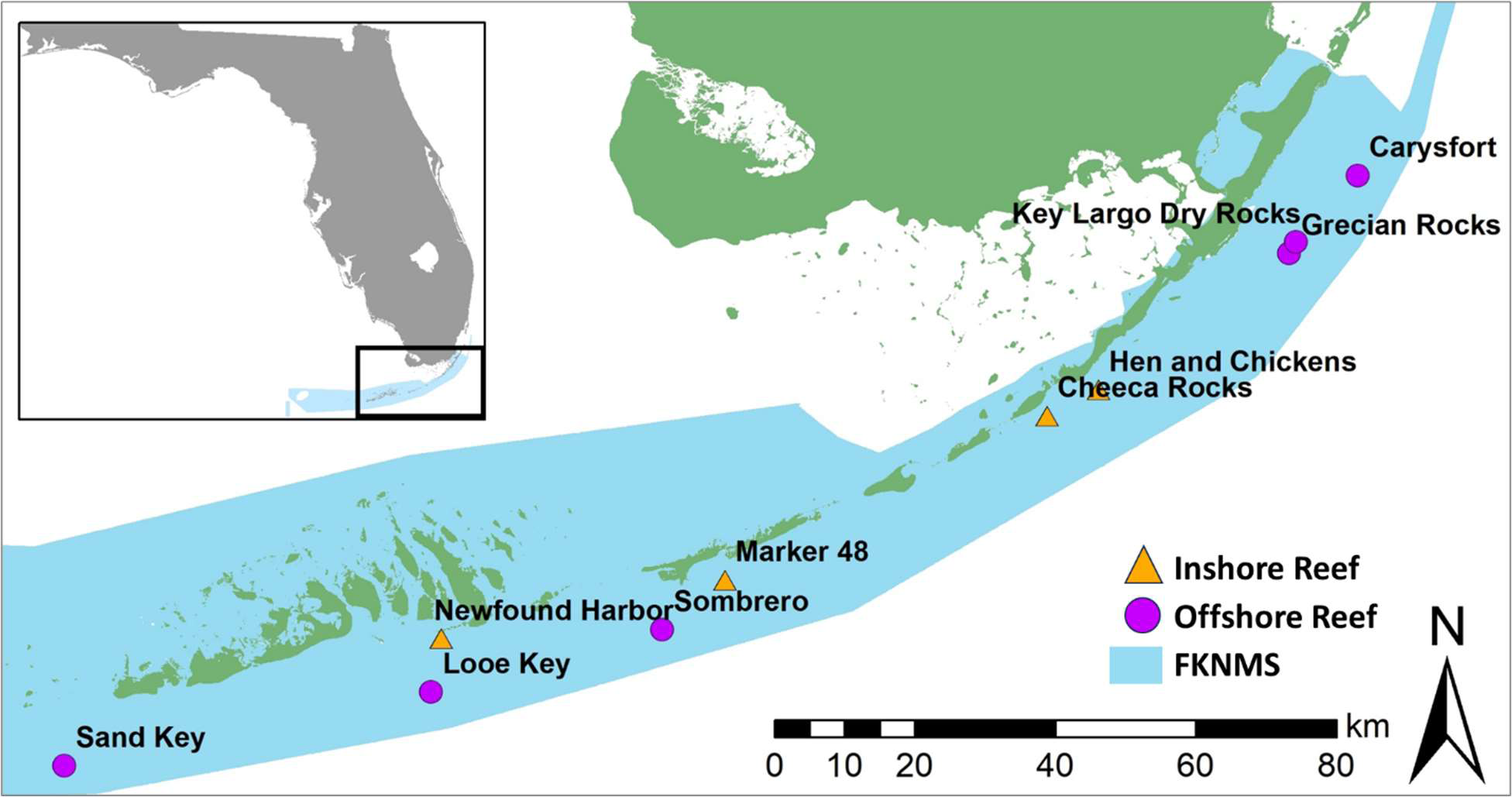
Location of regularly monitored reef sites in the Florida Keys. Four inshore sites (orange triangles) and six offshore sites (purple circles) are all located within the Florida Keys National Marine Sanctuary (FKNMS).

**Table 1.** Proportion of monitored corals exhibiting tissue loss from various stressors during each site visit. The length of the red bars increases with the proportion of the population affected.

|  | Site | Date | # corals | Max DHW since last visit | % with OF-unk | % with BBD | % with RM-BL | % DEAD-BL | % dead - unk |
| --- | --- | --- | --- | --- | --- | --- | --- | --- | --- |
| Offshore | Sand | 7/11-7/12/23 | 218 | 2.4 | 0.0% | 0.0% | 0.0% | 0.0% | 0.0% |
|  | Sand | 9/13/2023 | 218 | 16.3 | 0.0% | 0.5% | 0.0% | 0.0% | 0.0% |
|  | Sand | 11/27/2023 | 218 | 18.4 | 2.8% | 0.0% | 1.4% | 0.0% | 0.5% |
|  | Sand | 2/21-2/22/2024 | 217 | 2.9 | 1.8% | 0.0% | 0.0% | 0.0% | 1.4% |
|  | Looe | 6/27-7/7/2023 | 1148 | 2.2 | 0.0% | 0.0% | 0.0% | 0.0% | 0.1% |
|  | Looe | 9/5-9/7/2023 | 1147 | 16.3 | 0.0% | 0.0% | 0.0% | 0.0% | 0.0% |
|  | Looe | 11/22-12/1/2023 | 1147 | 18.0 | 12.6% | 3.2% | 0.5% | 0.0% | 1.4% |
|  | Looe | 1/31-2/25/2024 | 1131 | 2.6 | 0.7% | 0.3% | 0.0% | 0.0% | 1.2% |
|  | Sombrero | 7/13-7/14/2023 | 132 | 4.3 | 0.0% | 0.0% | 0.0% | 0.0% | 0.0% |
|  | Sombrero | 9/14/2023 | 132 | 17.4 | 0.0% | 0.8% | 0.0% | 0.0% | 0.8% |
|  | Sombrero | 12/5/2023 | 132 | 18.8 | 0.0% | 0.0% | 0.0% | 0.0% | 0.0% |
|  | Grecian/KLDR | 7/19/2023 | 99 | 4.3 | 0.0% | 0.0% | 0.0% | 0.0% | 0.0% |
|  | Grecian/KLDR | 9/20-9/21/2023 | 99 | 16.2 | 0.0% | 0.0% | 1.0% | 0.0% | 0.0% |
|  | Grecian/KLDR | 1/13/2024 | 96 | 16.2 | 0.0% | 0.0% | 0.0% | 0.0% | 4.2% |
|  | Carysfort | 7/20/2023 | 102 | 4.6 | 0.0% | 0.0% | 0.0% | 0.0% | 0.0% |
|  | Carysfort | 9/20-9/21/2023 | 102 | 16.2 | 0.0% | 0.0% | 0.0% | 0.0% | 0.0% |
|  | Carysfort | 1/18/2024 | 101 | 16.2 | 0.0% | 0.0% | 0.0% | 0.0% | 3.0% |
| Inshore | Newfound | 7/25-7/26/2023 | 547 | 11.9 | 0.0% | 1.1% | 11.0% | 6.2% | 0.0% |
|  | Newfound | 9/25-9/26/2023 | 513 | 21.1 | 0.0% | 2.3% | 17.0% | 10.1% | 2.7% |
|  | Newfound | 2/26-3/5/2024 | 447 | 19.7 | 0.0% | 0.2% | 0.0% | 0.0% | 28.6% |
|  | Marker 48 | 7/13/2023 | 357 | 4.1 | 0.0% | 0.0% | 0.0% | 0.0% | 0.6% |
|  | Marker 48 | 9/14-9/15/2023 | 355 | 18.0 | 0.0% | 0.3% | 0.8% | 0.3% | 0.8% |
|  | Marker 48 | 12/6/2023 | 351 | 19.2 | 0.0% | 0.0% | 0.0% | 0.0% | 0.3% |
|  | Marker 48 | 5/1-5/2/2024 | 350 | 2.1 | 0.0% | 0.3% | 0.0% | 0.0% | 2.6% |
|  | Cheeca | 6/5-6/22/2023 | 1335 | 0 | 0.0% | 0.0% | 0.0% | 0.0% | 2.5% |
|  | Cheeca | 8/8-8/24/2023 | 1299 | 15.6 | 0.0% | 7.0% | 11.9% | 6.8% | 2.8% |
|  | Cheeca | 9/10-9/11/2023 | 326 | 16.8 | 0.0% | 3.7% | 19.3% | 18.4% | 6.1% |
|  | Cheeca | 10/10-11/20/2023 | 1229 | 18.9 | 9.2% | 0.7% | 1.6% | 0.8% | 6.6% |
|  | Cheeca | 3/20-4/25/2024 | 1144 | 7.1 | 0.0% | 0.0% | 0.0% | 0.0% | 5.7% |
|  | Hen&Chickens | 7/31-8/1/2023 | 468 | 8.4 | 0.0% | 0.0% | 0.0% | 0.0% | 0.0% |
|  | Hen&Chickens | 9/28-9/29/2023 | 468 | 17.4 | 0.0% | 0.2% | 0.0% | 0.0% | 0.4% |
|  | Hen&Chickens | 3/7-3/18/2024 | 466 | 15.7 | 0.0% | 0.0% | 0.0% | 0.0% | 1.1% |

Corals at each site were part of a multi-year disease intervention project on colonies affected by SCTLD lesions. Beginning in 2019 for six of the sites, 2020 for two sites, and 2021 for the ninth site, corals with active SCTLD were tagged using numbered cattle tags, mapped, identified to species, and measured. Tagged corals were all treated with the amoxicillin+Base2b treatment which is largely effective at halting SCTLD lesions (Neely et al., 2020; Neely et al., 2021b; Shilling et al., 2021; Walker et al., 2021). Each site was revisited approximately every two months and surveyed for any corals showing signs of recent mortality. Untagged corals with newly-present SCTLD lesions were tagged, treated, and added to the monitoring program when encountered. Tagged corals with new SCTLD lesions would be re-treated as needed. Some corals at some sites had received treatments as early as two months prior to the onset of the bleaching mortality surveys, while others had not received treatments in up to 4.5 years.

During each monitoring period, previously-tagged corals with signs of recent mortality were assessed for the cause of mortality (Figure 3). During the bleaching period (July 2023 – October 2023), sources of mortality were:

1. SCTLD
2. BBD: recent partial mortality as a result of black band disease
3. RM-BL: recent partial mortality as a result of bleaching
4. RM-BL + BBD: recent partial mortality as a result of bleaching, as well as active black band disease lesions
5. OF-unk: an unusual condition seen only on *O. faveolata* corals as they regained zooxanthellae. Characterized by multiple “lesions” scattered throughout the colony, but most often originating from the tops or ridges of the colonies.
6. Dead-BL: colony completely dead as a result of bleaching-related mortality
7. Dead: coral died since the last monitoring period, but it was not possible to conclude what the cause of death was.

**Figure 3.**
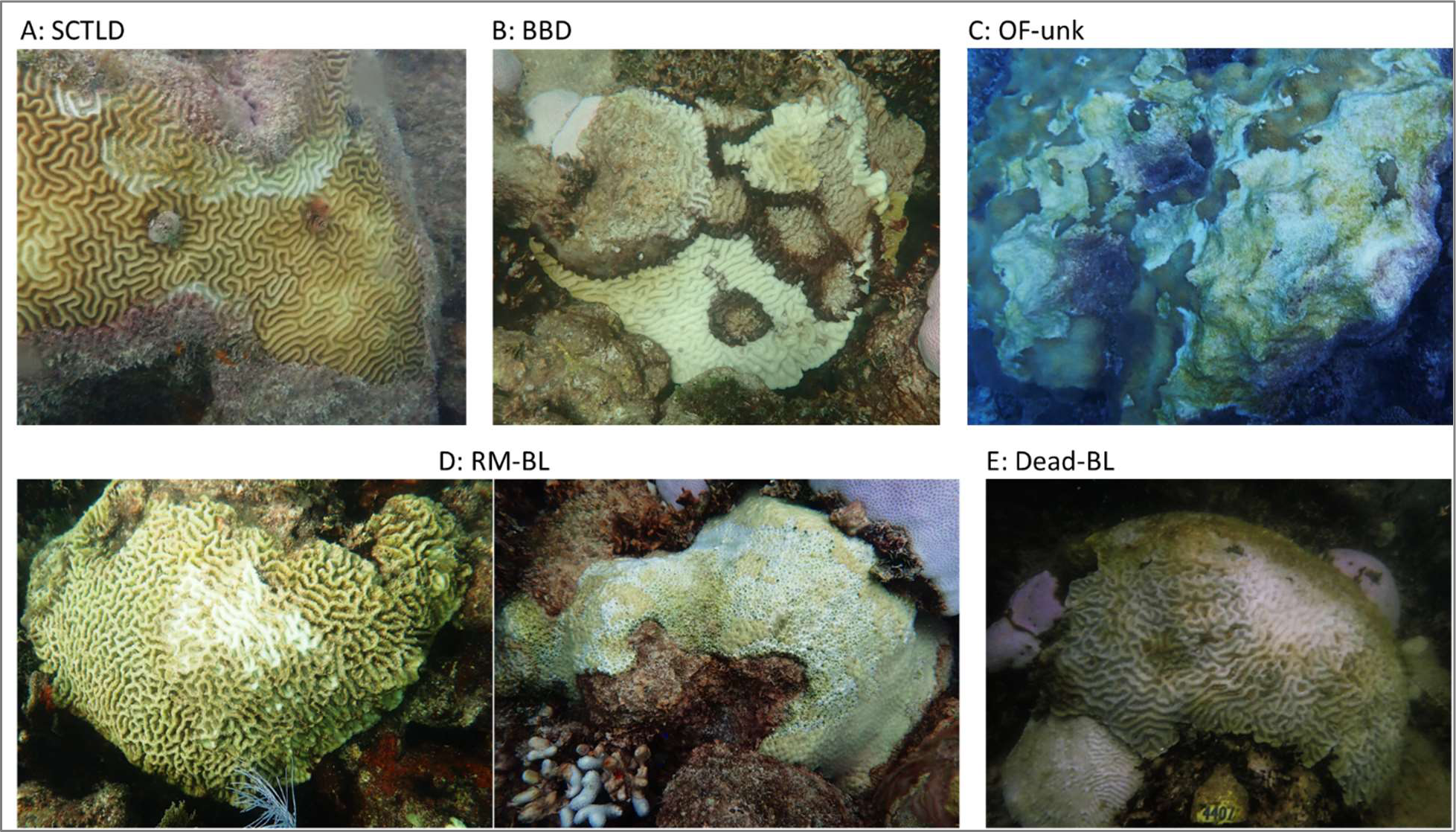
Images of the bleaching and post-bleaching sources of tissue loss observed on monitored corals. Stony coral tissue loss disease (SCTLD; A), black-band disease (BBD; B), *Orbicella faveolata* unknown condition (OF-unk; C), partial recent mortality from bleaching (RM-BL; D), and complete mortality from bleaching (Dead-BL; E)

Because each assessed coral was tagged and mapped, we were assured of monitoring the same corals over multiple time periods. Incidence data for each of the health status categories were based on a fixed number of corals and repeated over multiple time periods. However, because only colonies that had contracted SCTLD in the past were tagged, the monitoring excludes non-SCTLD susceptible species like the Acroporids and *Porites* branching species, as well as colonies of susceptible species that had not displayed active SCTLD lesions between the start of site monitoring and summer 2023. The relative abundance of species of monitored corals varied by site, as most offshore reefs were not initially incorporated into the SCTLD intervention program until many of the highly susceptible colonies had died off, while at inshore reefs, intervention was initiated earlier in the disease outbreak timeline, thus allowing for more of the highly susceptible species to survive (Figure 4).

**Figure 4.**
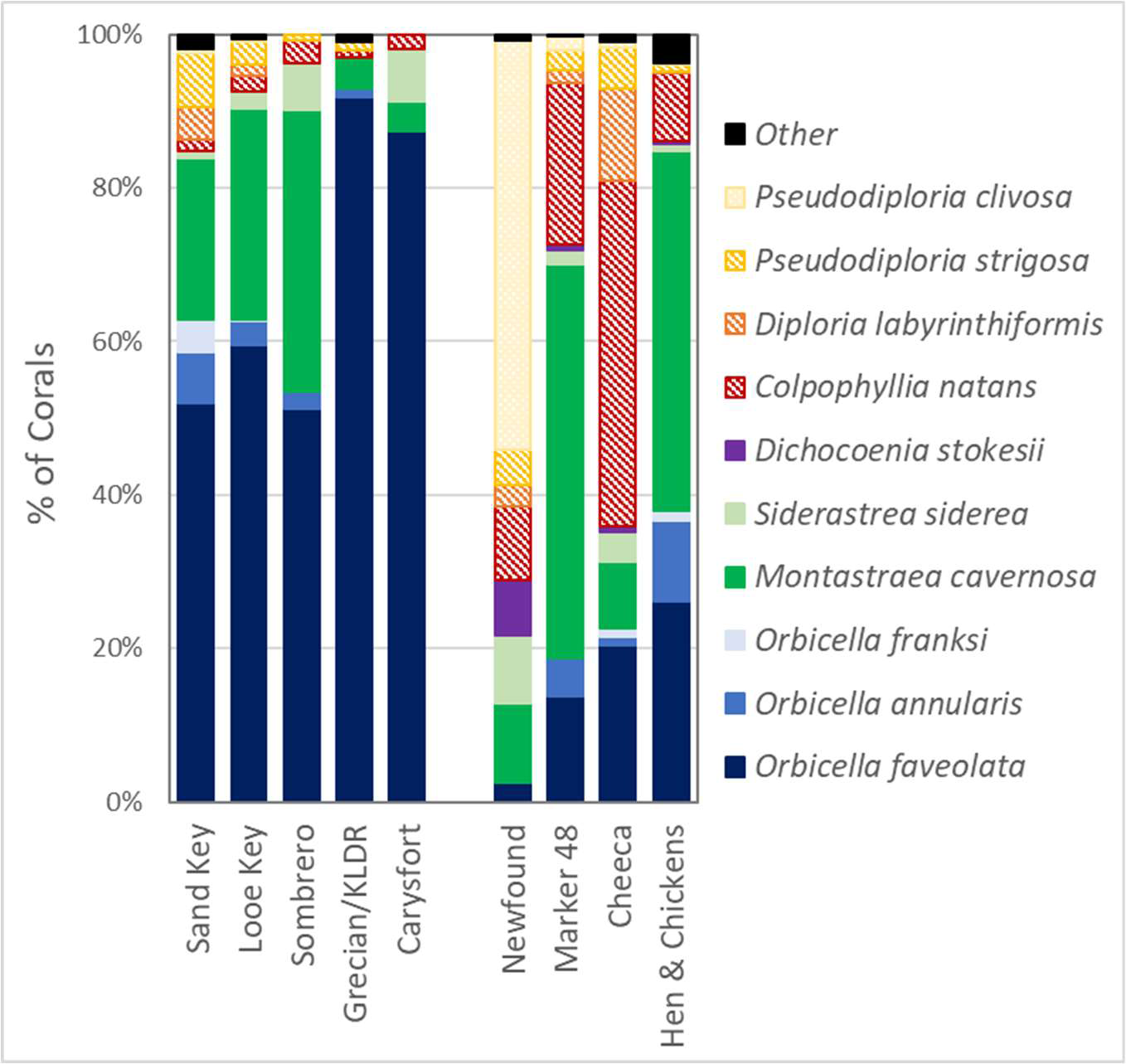
Proportion of monitored corals at each site by species. Boulder corals are solid cool tones, brain corals are crosshatched warm tones.

All reefs received post-bleaching assessments between December 2023 – April 2024 after thermal stress had subsided to pre-bleaching levels. At some of these reefs, unusual mortality patterns were seen on some *Orbicella faveolata* corals (Figure 3c). Lesions did not resemble those of traditional SCTLD, white plague, or other diseases. While these areas of recent mortality generally appeared on the top of coral colonies, particularly in the shallows (which may indicate a connection to the most thermally and UV-stressed areas), they were generally not on actively bleached tissue. Because the cause of mortality could not be determined, corals with this condition were coded “*O. faveolata* unknown” or OF-unk. We defined the proportion of *O. faveolata* colonies affected as the number of previously-tagged, live *O. faveolata* colonies with the condition, divided by the total number of previously-tagged, live *O. faveolata* colonies.

### Temperature Data

Sea surface temperature and cumulative thermal stress data were downloaded from NOAA’s Coral Reef Watch 5km single-pixel virtual station online products (NOAA Coral Reef Watch, 2023). These single-pixel stations were set up by NOAA specifically for the 2023 hyperthermal event, and strategically included our long-term monitoring stations (Figure 5). Sea surface temperatures are derived from satellite sensors and ground-truthed by operational buoys to maintain accuracy. Beginning in the summer of 2023, these data were shared daily in near-real-time.

**Figure 5.**
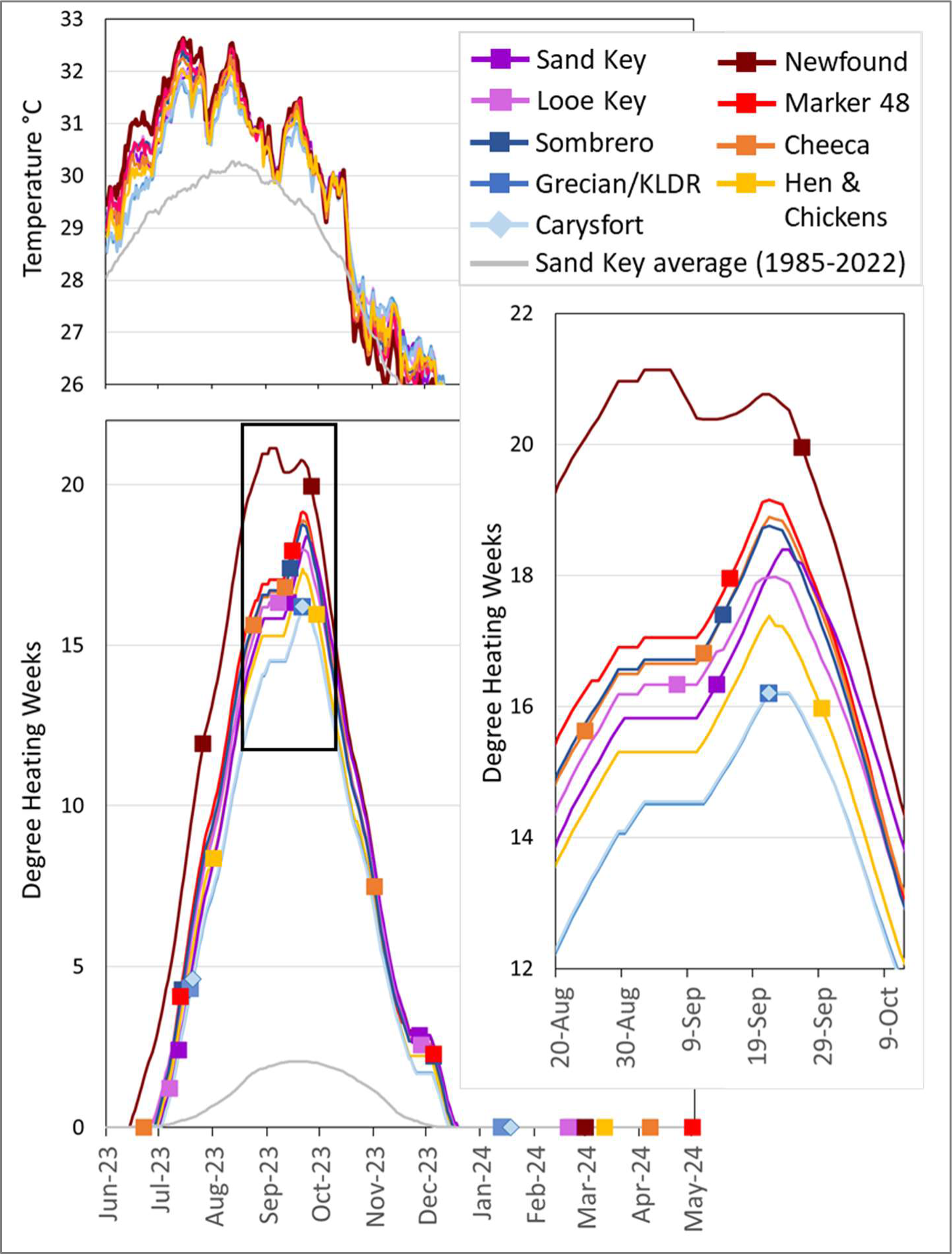
Temperature (top) and cumulative thermal stress (DHW; bottom) at monitored sites from June 1, 2023 through May 2, 2024. The average DHW and SST for Sand Key from 1985-2022 is displayed (grey line) for comparison. Squares on the cumulative thermal stress lines indicate the dates each site was monitored. For sites at which monitoring took multiple days, the mid-point of monitoring dates is indicated. The pop-out box (right) shows fine-scale differences between sites during peak thermal stress. Raw data are from NOAA’s Coral Reef Watch single-pixel virtual stations.

Degree heating weeks (DHW) are calculated using the sea surface temperature data. The metric combines the intensity of the heat stress (degrees C over the normal maximum) as well as duration (number of weeks of that stress), summed over the previous 12 weeks. DHW has been shown to be a more precise predictor of coral stress than temperature (Kayanne, 2017), with past accumulation of 4 DHW shown to cause significant coral bleaching, and values over 8 DHW known to cause severe bleaching and significant mortality (Kayanne, 2017).

### Analyses

#### Effect of site on mortality

To assess the effect of site on the risk of bleaching mortality for the three most common species (*Orbicella faveolata*, *Montastraea cavernosa*, and *Colpophyllia natans*), we performed a series of logistic regressions followed by pairwise site comparisons using the *stats* and *multcomp* packages in R (Hothorn et al., 2008; R Core Team, 2022). All pairwise comparisons had their p-values adjusted via the single-step method to correct for multiple comparisons. Prior to model construction, we binned partial– and full-bleaching mortality together to capture bleaching mortality holistically.

The first model for *O. faveolata* included all sites, including those with no mortality. Initial model results indicated high leverage for corals found at Newfound Harbor, so this site was dropped, and the model was run a second time. Significance of site as a global factor in mortality likelihood was assessed via chi-squared testing on the logistic regression model. Because site was found to be highly significant in both global models, Newfound Harbor was re-introduced for pairwise comparisons. Furthermore, because three sites (Carysfort, Hen and Chickens, and Sombrero) had no mortality and therefore no variability to model, they were removed prior to pairwise comparisons.

For the *M. cavernosa* model, the two sites with 20 or fewer corals were removed (Grecian/KLDR and Carysfort). The resulting dataset included seven sites: Sand Key, Looe Key, Sombrero, Newfound Harbor, Marker 48, Cheeca Rocks, and Hen and Chickens. The logistic regression indicated a significant effect of site, so follow-up pairwise comparisons were performed as outlined above.

For the *C. natans* model, the four sites with 20 or fewer corals were removed (Sand Key, Sombrero, Grecian/KLDR, and Carysfort Reef). The resulting dataset included five sites: Looe Key, Newfound Harbor, Marker 48, Cheeca Rocks, and Hen and Chickens. The logistic regression indicated a significant effect of site, so pairwise comparisons between the three sites with mortality – Cheeca Rocks, Marker 48, and Newfound Harbor – were performed.

#### Effects of species on mortality

To complement the species specific, cross-site comparisons, we also performed logistic regressions at the two sites where both species diversity and bleaching-related mortality were high (Newfound Harbor and Cheeca Rocks). At each site, we removed any species with fewer than 14 colonies represented. We then performed a global logistic regression to assess whether species had a significant effect on likelihood of bleaching mortality. Finally, as with the cross-site analyses, since the global model was significant, we performed pairwise comparisons to determine which species pairs were significantly different from one another, in which case holm correction factors were used to assist in model convergence with low sample sizes.

At the two high-mortality sites, we also assessed odds ratios of the species with at least 14 monitored colonies. We used the *epitools* package in R to measure the conditional maximum likelihood estimation via the Fisher method (Aragon et al., 2020). We used *C. natans* as the anchor species to normalize other results to, because of its prevalence and intermediate mortality rate at both sites.

#### Effect of pre-bleaching history

At Cheeca Rocks, a large number of corals exhibited SCTLD lesions and were treated with topical amoxicillin in the months immediately preceding the hyperthermal event. This provided an opportunity to assess how these diseased/treated corals fared through the event in relation to non-diseased/non-treated corals. For the four common species in which bleaching-related mortality was observed – *P. strigosa*, D*. labyrinthiformis*, *C. natans,* and *O. faveolata* – we compared the numbers of corals exhibiting bleaching-related mortality (either full (Dead-BL) or partial (RM-BL)) for corals that were healthy in June to those that were diseased/treated in June. We compared the treatment groups using chi-squared tests with a Bonferroni correction (α = .0125).

We did a similar analysis to assess whether pre-bleaching health status in June 2023 correlated with the appearance of black band disease in August 2023. Six common species developed black band disease – *P. strigosa*, *D. labyrinthiformis*, *C. natans*, *O. faveolata*, *M. cavernosa*, and *S. siderea*. For each, we compared the number of corals with black band disease and the number of living corals without black band (any corals that were dead in August 2023 were excluded from analyses) between the two June 2023 health groups. We used chi-square tests with a Bonferroni correction (α = .0083).

## Results

### Summary of impacts

Across all offshore sites, bleaching-related mortality and black band disease prevalence were both very low in spite of 100% bleaching (Table 1). Of the 1635 offshore monitored colonies, only one, an *O. faveolata* at Grecian Rocks, had any partial mortality related to bleaching. Only two colonies, a *M. cavernosa* at Sand Key and a *M. cavernosa* at Sombrero, had black band disease.

At inshore sites (all of which also had 100% bleaching), mortality relating to the MHW was more variable. Two sites, Marker 48 and Hen & Chickens, had minimal mortality. At Marker 48, there was no bleaching-related mortality or black band disease in July, and five colonies with partial or full mortality in September. Overall, of the 348 colonies monitored at this Middle Keys reef, 0.3% had black band, 0.9% had partial mortality from bleaching, and 0.3% died from bleaching. At Hen and Chickens, values were similarly low, with no black band or bleaching-related mortality observed in early August and only a single colony with black band disease in late September.

At the two other inshore reefs, losses were greater. Newfound Harbor was already experiencing thermal-related losses during the first visit in July (11.9 DHW). Of the 547 monitored colonies, 0.2% were afflicted with black band disease, 1.0% had both black band and bleaching-related mortality, 10.7% had partial mortality due to bleaching, and 6.2% were already fully dead as a result of bleaching. Two months later (513 colonies monitored), black band incidence had increased to 1.9%, while 0.4% of colonies had both black band and bleaching-related mortality, 15.8% had partial mortality due to bleaching, and an additional 10.1% had died completely as a result of bleaching. An additional 2.7% of colonies had died since the previous visit but their cause of death could not be confirmed. By March 2024 (447 remaining colonies monitored), bleaching-related partial mortality was non-existent, and black band disease was present on only a single colony. However, an additional 29% of colonies had died since the previous monitoring event. In total, 42.9% of monitored colonies at Newfound Harbor perished completely during the hyperthermal event, and an additional 12.4% exhibited partial mortality from bleaching and/or black band disease.

Colonies at the inshore Cheeca Rocks site were also heavily impacted. During August (15.6 DHW), incidence of various stressors on the 1299 monitored colonies was: 5.7% black band disease, 1.3% black band disease as well as bleaching-related partial mortality, 10.5% bleaching-related partial mortality, 6.8% full mortality from bleaching, and 2.8% full mortality from indeterminate cause. One month later, within the subset of 326 colonies, black band incidence had declined to 2.0%, and the remaining lesions had smaller more diffuse disease mats and slower progression rates. However, the proportion of colonies with recent mortality related to bleaching had increased to 13.1%. An additional 4.4% of colonies had died completely as a result of bleaching, and an additional 2.2% had died without a determinant cause. By October/November, black band disease incidence was 0.7%, 1.6% of colonies had active partial mortality from bleaching, 0.8% were recently dead from bleaching, and 6.6% were dead from indeterminate cause. When colonies were revisited in April 2024, there was no black band disease or bleaching-related mortality, but an additional 5.7% of colonies had died since the previous monitoring. Overall, at Cheeca Rocks, 24% of corals experienced complete mortality through the bleaching event while an additional 14% had partial mortality due to bleaching and/or black band disease.

### Effect of site on mortality

The regression models identified some significant site-specific differences in mortality (Figure 6). For both global logistic regression models (including and excluding Newfound Harbor) for *O. faveolata*, site had a strongly significant effect on likelihood of mortality (Analysis of deviance. p < 0.0001 for both), indicating that the effect of site was not driven only by high mortality at Newfound Harbor. From a pairwise perspective, Newfound Harbor had significantly higher mortality than all tested sites (Tukey tests. Sand Key p = 0.02; Looe Key p = 0.005; Grecian/KLDR p = 0.02) except Marker 48 (which had a small sample size) and Cheeca Rocks, which had the second highest mortality rate. While mortality at Cheeca Rocks was significantly higher than only Looe Key in the model (Tukey test p = 0.006), the fact that all other sites aside from Newfound Harbor had lower mortality than Looe Key suggests that *O. faveolata* at Cheeca Rocks had a higher mortality risk than all sites except Newfound Harbor.

**Figure 6.**
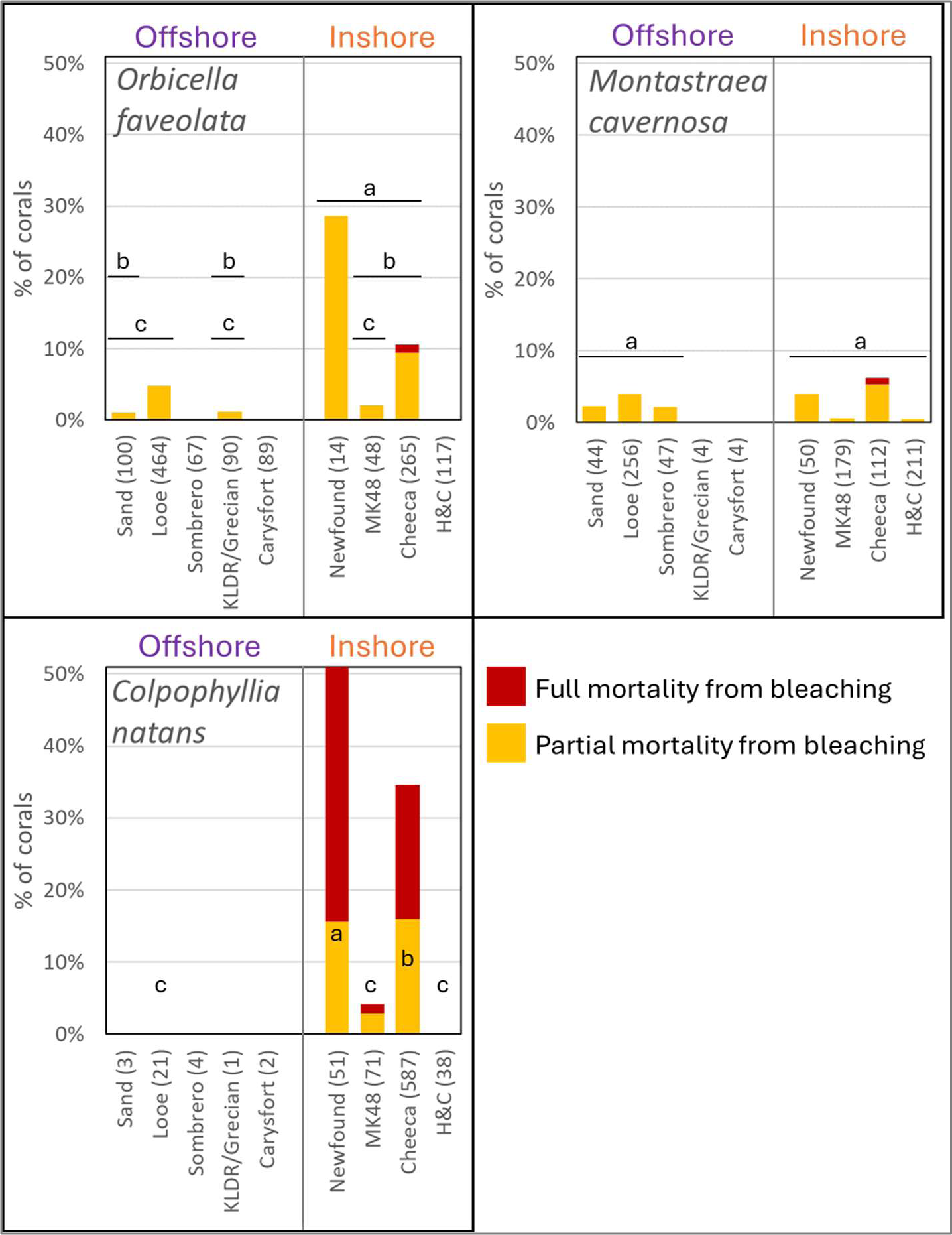
Proportion of monitored corals exhibiting partial (yellow) or full (red) bleaching-related mortality. Sites are separated by offshore/inshore and displayed west to east. The number of colonies assessed in the logistic regression models for each species is shown after the site name. Within each species, sites not significantly different from each other share a lettered bar. Sites with fewer than 20 corals of a species were excluded from the models, and sites with no mortality were excluded from pairwise comparisons; for these, no significance letters are displayed.

Site had a moderately significant effect on *M. cavernosa* bleaching-related mortality (Analysis of deviance. p = 0.015). However, corrected pairwise comparisons identified no significant differences between reefs. This discrepancy is likely driven by the multiple comparison penalty corrections imposed in the pairwise model. As a result, it is likely that actual site level differences are real, but minimal for this species.

Site had a highly significant effect on *C. natans* bleaching mortality (Analysis of deviance. p < 0.0001), driven by high mortality at Newfound Harbor and Cheeca Rocks, minimal mortality at Marker 48, and no mortality at other sites. Newfound Harbor had the greatest mortality, significantly higher than at Cheeca Rocks (p = 0.049) and Marker 48 (p < 0.001) and by extension Hen and Chickens and Looe Key. Cheeca had greater mortality than Marker 48 (p < 0.001), and by extension Hen and Chickens and Looe Key.

### Fate tracking

For the three sites with the highest bleaching-related mortality – Newfound Harbor, Marker 48, and Cheeca Rocks – we assessed how the status of individual corals changed through multiple months of the bleaching event (Figure 7). At Newfound Harbor, 81% of monitored corals had no signs of mortality in July. Of those, 70% continued to show no signs of mortality through September, while 21% developed bleaching-related tissue loss, and 10% had died by September. Conversely, 11% of corals had bleaching-related mortality in July; 40% of these “recovered” to a status of no active tissue loss, 19% continued to show bleaching-related partial mortality, and 41% died. Though nearly double the proportion of corals transitioned from partial-mortality to no-mortality than vice versa, death and partial mortality were affecting more corals in September as a result of the large number of previously “healthy” corals that transitioned to partial and total mortality. The corals continued to experience mortality between the September 2023 and March 2024 monitoring events. Though 70% of the black band diseased corals and 4% of the corals with bleaching-related partial mortality in September recovered to non-mortality status, 56% of corals with partial mortality as well as 24% of corals showing no signs of mortality in September were dead by March 2024.

**Figure 7.**
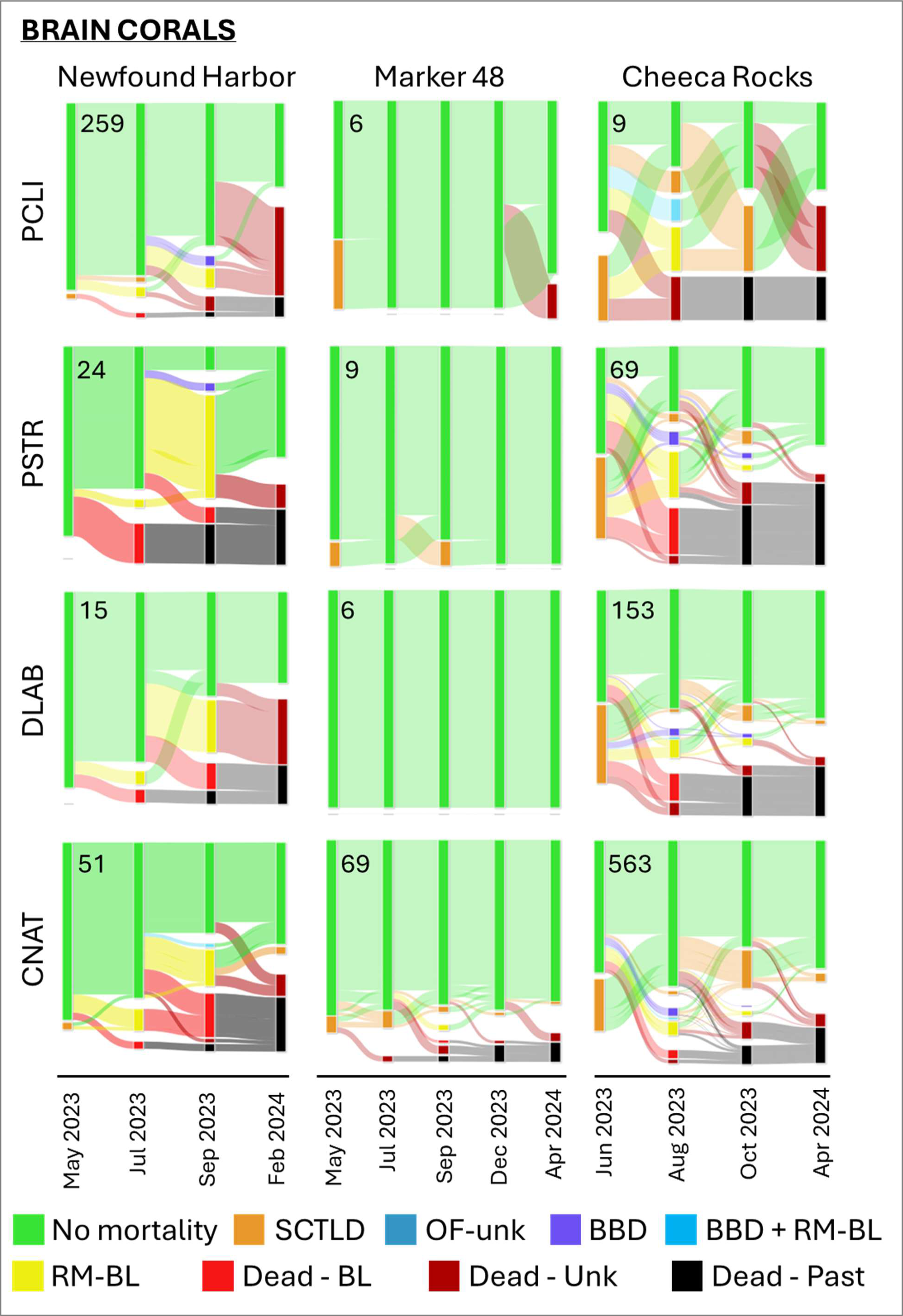
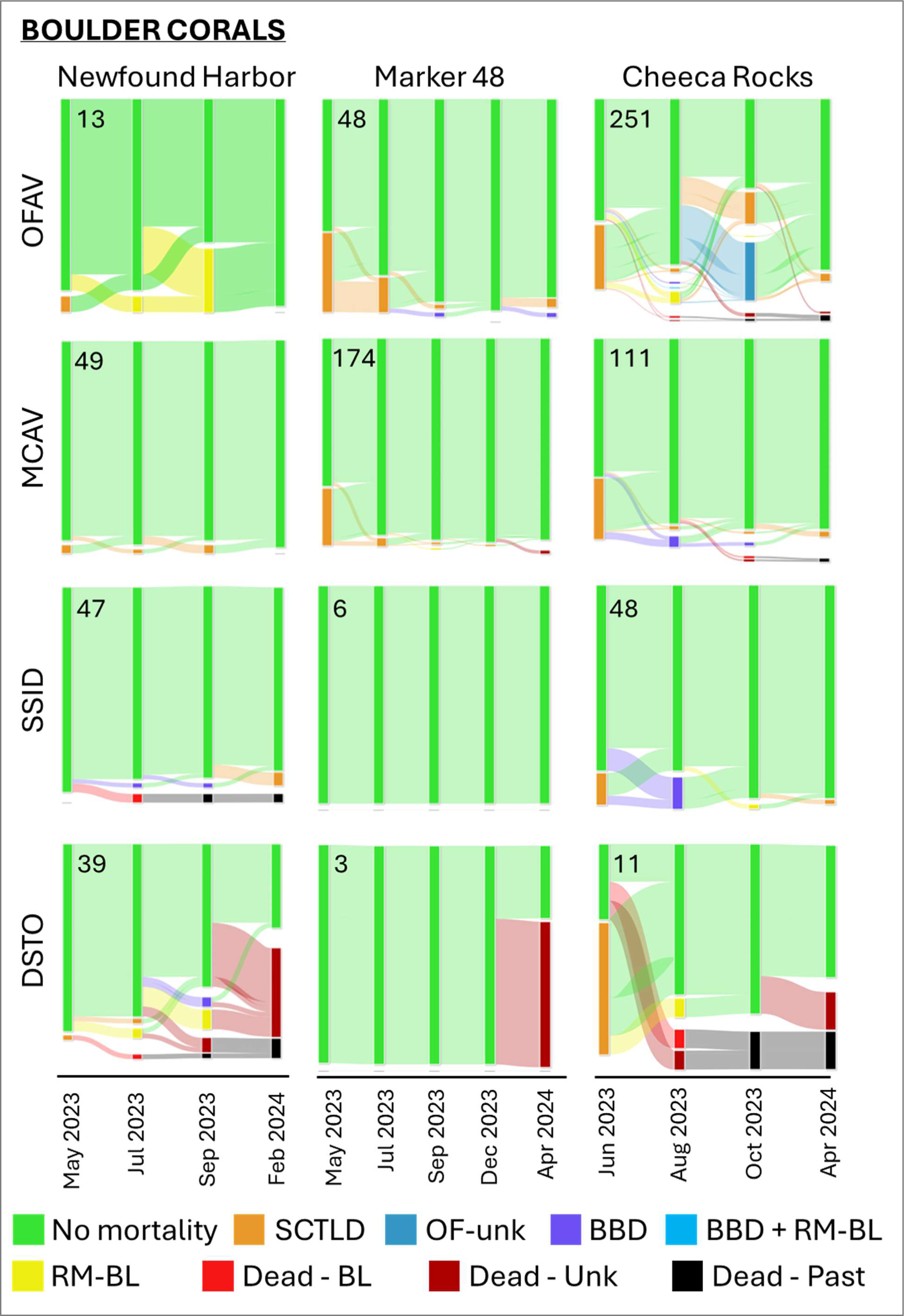
The proportion of fate-tracked corals within each health status across time periods at three inshore reefs, and the movement of corals between health categories across monitoring events. Sample size for each is shown in each plot.

At Marker 48, there were no signs of thermal-related mortality in July. Of the four colonies that exhibited partial mortality in September, all had recovered by December.

The subset of corals monitored monthly at Cheeca Rocks was also assessed for individual colonies’ status through time. The proportion of colonies showing no signs of mortality was at its lowest during the September and October assessments (54%). Black band disease was highest in August and continued to decline through subsequent months. Bleaching-related partial mortality was 14-15% during both August and September, but declined to 4% in October and was 0% by March 2024. Between August and September, 13% of corals without signs of mortality developed partial bleaching-related mortality, while 20% of those with partial mortality recovered to a no-mortality state. From September to October, only 2% of non-affected corals transitioned to a bleaching-related mortality state, while 41% of those with bleaching-related mortality recovered to a no-mortality state. By March 2024, an additional 7% of corals had died. These newly dead corals were mostly from those with partial mortality in October 2023: 35% of corals with bleaching-related partial mortality, 21% of those with treated SCTLD lesions, and 6% of those with no signs of mortality in October. On the recovery side, 30% of those with treated SCTLD lesions and 64% of corals with partial mortality from bleaching in October appeared fully healthy in March. Though stony coral tissue loss disease was prevalent at this site before the bleaching event, it declined to 1% in August, but began to reappear in subsequent months, up to 4% in September and 12% in October.

### Effect of species

At both Newfound Harbor and Cheeca Rocks, mortality varied significantly by species (p < 0.0001), with complex pairwise comparisons (Figure 8a). At both sites, *P. strigosa* colonies experienced the greatest mortality, with 88% of colonies at Newfound Harbor and 57% of colonies at Cheeca Rocks losing some or all of their tissue to bleaching. At both sites, the next most highly impacted species were the other brain coral species, while boulder corals had lower rates of mortality. At both sites, *M. cavernosa* was the least impacted species.

**Figure 8.**
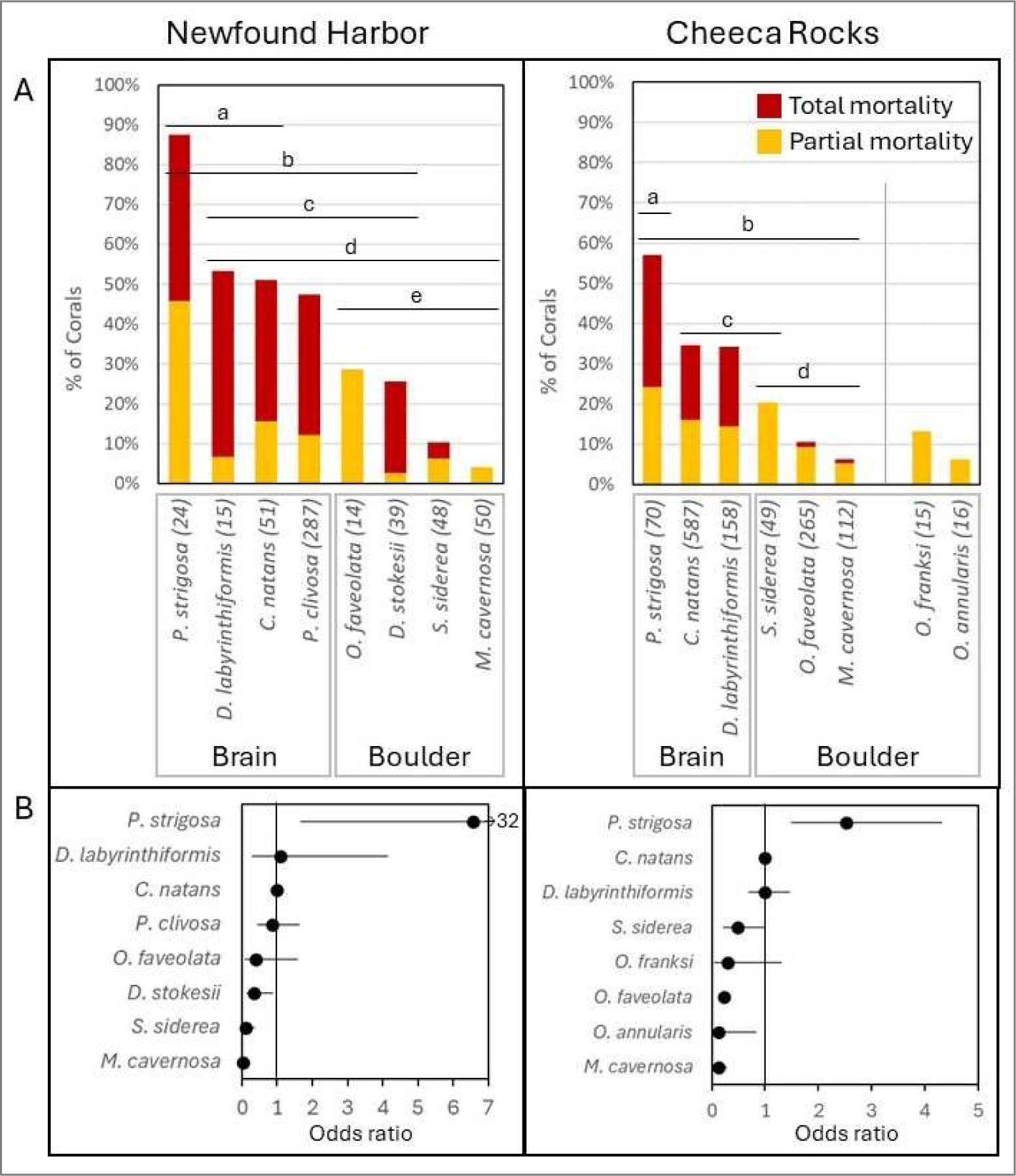
A) The proportion of colonies of various coral species exhibiting partial bleaching-related mortality (yellow) or full bleaching-related mortality (red) during the 2023 thermal bleaching event at the two most heavily impacted sites. Groups not significantly different from each other share a lettered bar. At Cheeca Rocks, *Orbicella franksi* and *O. annularis* had small sample sizes and did not differ significantly from any other species. B) Odds ratios for Newfound Harbor and Cheeca Rocks identify the odds of a coral species having higher or lower mortality in relation to the anchor species, *Colpophyllia natans*. Black dots indicate the odds ratio, with whiskers identifying the 95% confidence intervals. Whiskers that cross 1 do not have significantly different odds of mortality than *C. natans*.

Odds ratios identified the same patterns of mortality (Figure 8b). We used *C. natans* as the anchor species, and at both sites the same suite of species had higher and lower odds of mortality from that species. The brain coral *P. strigosa* was 6.6 times more likely to experience mortality than *C. natans* at Newfound Harbor, and 2.5 times more likely at Cheeca Rocks. The other brain corals had similar mortality odds ratios to *C. natans*; *D. labyrinthiformis* overlapped at both sites, and *P. clivosa* which was only abundant at Newfound Harbor overlapped at that site. Most boulder corals had lower odds of experiencing mortality than *C. natans*. At Newfound Harbor, the odds of mortality as compared to the *C. natans* anchor species decreased by 96% for *M. cavernosa*, 89% for *S. siderea*, and 66% for *D. stokesii*. The odds of mortality for *O. faveolata* decreased by 61%, but this was not significantly different. At Cheeca Rocks, the odds of mortality compared to *C. natans* decreased by 87% for *M. cavernosa*, 87% for *O. annularis*, and 88% for *O. faveolata*. Odds decreased by 81% for *O. franksi* and 51% for *S. siderea*, but these were not significantly different.

### Effect of pre-bleaching history

We assessed the status of corals from six species in August, comparing individuals that were healthy in June to those which had SCTLD lesions and were treated (Figure 9). For four of the six species, black band incidence was higher on corals that had SCTLD lesions in June, but only for *C. natans* was this difference significant (p < 0.007; α = 0.0083). For two species, the proportion of corals exhibiting bleaching-related mortality (partial or total) was greater for corals that had SCTLD in June than for those that had been healthy, but this difference was only significant for *C. natans* (p < 0.001; α = 0.0125).

**Figure 9.**
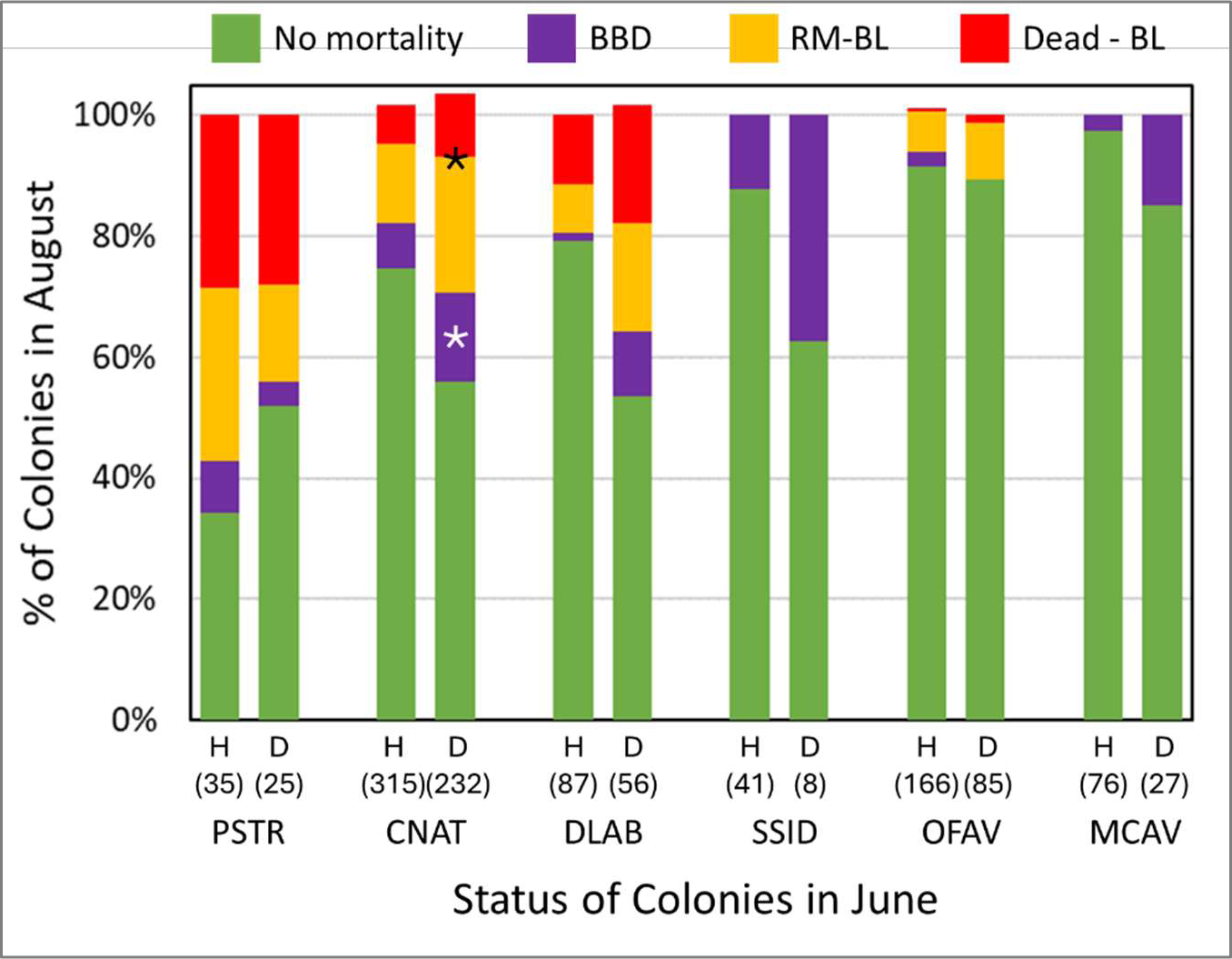
The proportion of corals exhibiting different health statuses at Cheeca Rocks in August: no mortality, black band disease (BBD), bleaching-related partial mortality (RM-BL), and dead from bleaching (Dead-BL). Corals are separated by species, and by their health status in June – healthy (H) or diseased and treated for SCTLD lesions (D). Species are represented by four letter codes: PSTR = *Pseudodiploria strigosa,* CNAT = *Colpophyllia natans*, DLAB = *Diploria labyrinthiformis*, SSID = *Siderastrea siderea,* OFAV = *Orbicella faveolata, and* MCAV = *Montastraea cavernosa.* Values do not always sum to 100% because a small number of colonies exhibited both RM-BL and BBD. Significant differences between the two groups within each species are indicated by *.

### Post-bleaching *O. faveolata* mortality

Though most sources of mortality abated upon coral color returning, some sites experienced an unknown source of mortality on *O. faveolata* colonies. Prevalence of the “OF-unk” condition varied greatly. At the offshore sites, 6% (6 of 109) of tagged *O. faveolata* colonies at Sand Key, 19% (121/654) at Looe Key, and 0% at Sombrero (0 of 66), Grecian/KLDR (0 of 90), and Carysfort (0 of 89) exhibited this condition. At inshore sites, none of the *O. faveolata* at Newfound Harbor (0 of 14), Marker 48 (0 of 48), or Hen and Chickens (0 of 117) exhibited signs of OF-unk, but 35% (87 of 251) of *O. faveolata* at Cheeca Rocks did. At the two heavily affected sites (Looe Key and Cheeca Rocks), we compared the health status of the corals before bleaching to assess whether that influenced their likelihood of developing OF-unk lesions after bleaching, but found no differences. At Looe Key, 19% of OF-unk colonies (26/135) were SCTLD-infected in June, compared to 18% (95/519) of non OF-unk affected colonies (chi-squared: p = 0.9). At Cheeca Rocks, 36% (29/80) of OF-unk affected colonies had SCTLD in June, compared to 34% (58/171) of non-OF-unk colonies (chi-squared: p= 0.8).

## Discussion

Fate-tracked brain and boulder coral monitoring through the 2023 MHW that caused near 100% bleaching in the Florida Keys identified extremely low mortality rates overall, but also some spatial and species-specific patterns. In general, offshore corals fared well through the bleaching event, with no colony-level losses to bleaching or black band disease. In contrast, some inshore sites fared worse, though losses were not uniform across inshore reefs. These spatial patterns largely match temperature and cumulative thermal stress variation among the reefs. Overall, offshore and more northern (Upper Keys) waters remained cooler and had less cumulative thermal stress than inshore and more southern (Lower Keys) waters (Figure 5). With the exception of Hen and Chickens, all inshore reefs experienced greater cumulative thermal stress than all offshore reefs. This is not surprising, as inshore waters are generally shallower and more affected by the shallow Florida Bay waters moving between islands, while offshore reefs are generally slightly deeper with temperatures more moderated by the proximity to the Gulf Stream. However, the cooler waters at Hen and Chickens reef compared to Cheeca Rocks, which is only 7.5 km away and approximately the same distance from shore, is surprising. This temperature difference is consistent with the vastly different outcomes in coral survival between the two reefs.

In general, corals on Florida Keys inshore reefs now pale or bleach during most summers as a result of warm water temperatures. It is possible that as a result, the corals that remain are more resistant or resilient to thermal stress and potentially stress-hardened, though whether the driver of this is underlying genetics or some induced process is unclear. Though they certainly have fared better through past bleaching events than corals at offshore sites, acute thermal stress from the 2023 unprecedented MHW was too extreme for colonies at two of these sites. Notably, however, the extensive loss of corals at Newfound Harbor was observed as early as the July monitoring event when DHW were 11.9 – a level of cumulative thermal stress far below what was present study-wide later in the summer. We suggest that at this reef, losses were a direct result of abnormally high acute temperatures rather than accumulated thermal stress; the losses were not due to “bleaching” in the sense that corals were dying of depleted energy reserves from zooxanthellae loss, but instead a result of exceeding their thermal maxima and dying from hyperthermia. To our knowledge, this condition has not been previously documented in wild corals (but see Leggat et al. (2019) for similar observations in laboratory experiments).

Though it is tempting to assume that corals at sites faring well or poorly through the 2023 event may have certain resiliency or susceptibility, it is important to note that conditions within a reef and within a region experience high variability in weather patterns, changes in current as a result of variations in global circulation, waterflow from upstream, wind, and tidal influences, variation in cloud cover and turbidity, and other factors. There is no guarantee that sites will perform the same in future years, and we caution against making management decisions such as outplanting regimes or spawning hubs based on one year’s worth of temperature and survival data. We did, however, use the Coral Reef Watch single-pixel database to track cumulative thermal stress (DHWs) back to 1985. We identified the maximum DHW value for each year at each site. For the 14 years in which the maximum DHW value at any site exceeded 4, we ranked each of the sites against each other in terms of thermal stress (Figure 10). The warmest of the 9 sites received a value of 1, and the coolest a value of 9. We found that while sites were not consistent across all years in their cumulative thermal stress rankings compared to other sites, there were patterns. Newfound Harbor was consistently one of the most thermally stressed reefs, while Grecian/KLDR and Carysfort were consistently the coolest. Hen and Chickens consistently ranked lower for thermal stress than other inshore reefs as well as most offshore reefs. These temporal patterns suggest that some sites, even sometimes those in close proximity, may have oceanographic or bathymetric features that promote patterns of greater or lesser thermal stress than other reefs in the region. These should be further explored as possible factors for management actions.

**Figure 10.**
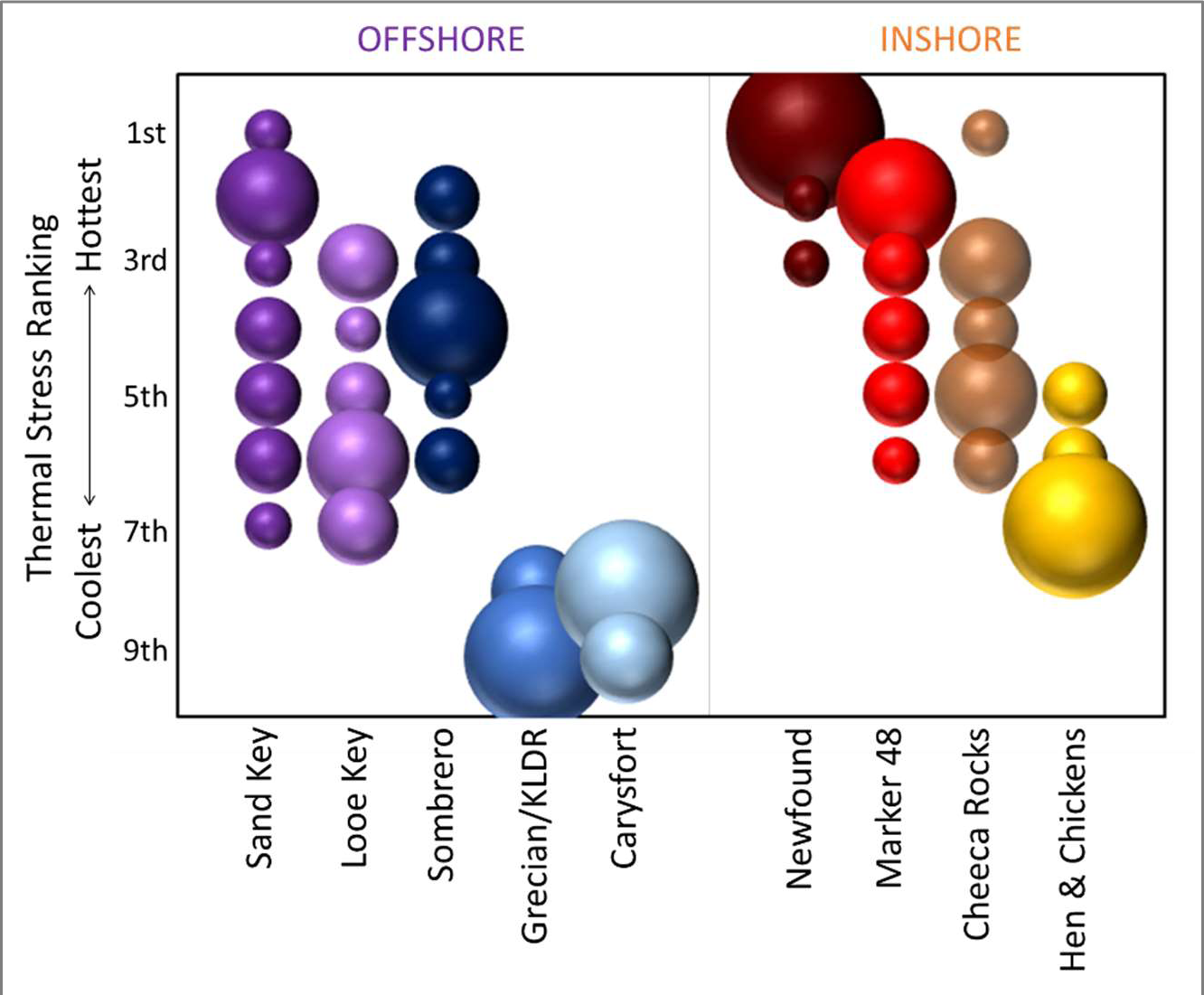
The comparative ranking of maximum cumulative thermal stress values across the nine study reefs for years 1985-2023 in which degree heating weeks (DHW) exceeded 4 for at least one site. The size of the bubble indicates the number of years a site placed in that ranking.

This history of differential thermal stress may also provide a potential explanation for the substantial losses of corals at Cheeca Rocks as compared to Marker 48, even when losses within individual species are compared. Though Marker 48 experienced greater DHWs in 2023 than Cheeca Rocks, coral mortality at Cheeca Rocks was much greater. We propose three possible hypotheses.

1) Marker 48 has repeatedly experienced higher thermal stress than Cheeca in past years, including a peak of 7.3 DHW in 2007 while Cheeca has never experienced DHWs greater than 6.3. While these cumulative thermal stress levels are minor in comparison to the values of 19 DHW that both sites experienced in 2023, it is possible that greater thermal stress at Marker 48 in the past had either killed off thermally intolerant individuals or helped to stress-harden the survivors.
2) Though both sites experience variable water quality and are susceptible to high turbidity from nearshore runoff and wave action, Marker 48 is, in general, a more turbid site compared to Cheeca Rocks. This may have resulted in reduced UV radiation, which may have minimized bleaching impacts (similar to results by Sully and van Woesik (2020)).
3) Though we generally reject the idea of site resilience due to the correlations of higher bleaching-related mortality with greater DHW levels, this comparison between Marker 48 and Cheeca does defy that trend. It is possible that colonies at Marker 48 do have some sort of resilience that enabled them to survive the same temperatures and cumulative thermal stress conditions that resulted in substantial mortality at Cheeca Rocks. Transplant experiments or laboratory testing of corals from these two sites may provide insight into this possibility.

Though coral fates among sites are largely explained by variation in cumulative thermal stress, we show conclusively that some species perform better than others under the same thermal conditions. At both sites experiencing substantial mortality, brain corals had higher mortality rates than boulder corals. *Pseudodiploria strigosa* colonies in particular were highly susceptible to bleaching-related mortality. To our knowledge, this is the first observation of differential mortality rates among these two coral growth forms. However, Brandt (2009) found similar patterns in non-lethal bleaching extent and bleaching duration during the 2005 Florida hyperthermal event, with *C. natans* and *P. strigosa* remaining bleached for longer and having greater proportions of their tissue area bleached than *M. cavernosa*, *O. faveolata*, and *S. siderea*. Loya et al. (2001) suggested that different survival rates between branching and massive corals following a bleaching event in Japan could be due to greater tissue thickness among the “winners”, which could offer self-shading to underlying zooxanthellae. However, patterns of tissue thickness do not correlate with our observed patterns of mortality, as the thick-tissued *C. natans* and *P. strigosa* (Smith et al., 2013) were particularly susceptible to mortality while the thin-tissued *S. siderea* was not. The role of symbionts in bleaching susceptibility is also widely known, with some species known to be more thermally tolerant than others. While the corals monitored in this study were not specifically sampled for symbiont identification, other studies have not documented widespread differences in symbiont communities between brain and boulder corals; the species assessed here contain predominantly *Breviolum* or *Cladocopium* (LaJeunesse, 2002; Correa et al., 2009; Grajales and Sánchez, 2016). As such, it is unlikely that genus-level symbiont differences are driving these differential patterns of mortality between brain and boulder corals. Alternate explanations for differential survival that may warrant future study include differences among species in energy reserves, degree of autotrophy, mass transfer efficiency (Loya et al., 2001), or other parameters.

Because the monitoring used to assess the trends in this paper was focused on SCTLD-affected corals, it excluded coral species that are seemingly immune to SCTLD such as the branching *Acropora* species. However, the authors anecdotally saw extremely high bleaching-related mortality rates of both wild and outplanted *Acropora palmata* and *Acropora cervicornis* colonies across the monitoring sites. There are two long-term *A. palmata* monitoring programs within the Florida Keys, both of which documented significant bleaching-related mortality following the 2014-2015 back-to-back hyperthermal events (Williams et al., 2017; Neely et al., 2022), and these programs may be expected to yield information in the future about the fate of remaining fate-tracked colonies through the 2023 event. A NOAA survey of *A. palmata* founder genotypes across Florida’s Coral Reef following the 2023 MHW documented extinction of 77% of genotypes in the wild (Williams, 2024). *Acropora cervicornis* monitoring in the Florida Keys is primarily via restoration programs. Early news reports and preliminary presentations suggest *A. cervicornis* mortality on both wild and outplanted colonies was extremely high, and it is hoped that future publications will be able to quantify these losses, as our observations suggest the Acroporids fared much more poorly than the SCTLD-susceptible species. Similar observations of branching corals faring more poorly than massive/boulder corals during other non-Caribbean bleaching events were reported by both Loya et al. (2001) and McClanahan (2004). While *A. palmata* and *A. cervicornis* were once dominant components of Florida Keys reefs, their widespread losses decades ago made them minor components of remaining reefs, with only a few wild thickets of each species remaining in the region before this event. Though the loss of wild individuals represents a blow to species diversity in the region, it has little impact on reef function or even coral cover and species diversity outside of those few small areas. On the other hand, it represents a devastating blow to restoration efforts in Florida which, though diversifying, still focus primarily on these two species. The increasing frequency and intensity of marine heatwaves may represent a call for restoration practitioners to assess the value of focusing primarily on species which seem so poorly suited for handling the climate of the immediate future. Acroporids appear very poorly suited for future climate conditions. But this study also shows that as temperatures continue to rise, and bleaching events become more severe, Caribbean restoration practitioners may also need to prioritize boulder corals over brain corals. In particular, *M. cavernosa* fared best at the highly impacted reefs, while *P. strigosa* colonies fared worst.

Even after MHWs subside, they can leave behind substantial legacies in a variety of marine benthic communities (summary in Smith et al. (2023). Though we show here that most sites experienced negligible losses to brain and boulder corals, and that even at the most heavily-impacted sites, most corals survived, the longer-term impacts of the bleaching event remain to be seen. Previous bleaching events have shown that bleached corals are more susceptible to diseases such as black band disease and white plague (Spadafore et al., 2021), presumably because stressed corals are immuno-compromised. Though we observed black band disease at affected sites peaking with water temperatures and then quickly subsiding even before color was returning to many corals, our observations of unusual lesions on *O. faveolata* at some sites as color returned to coral tissues suggests a level of bleaching impact which caused, in some cases, substantial mortality on survivors. In contrast, as seen in previous years in Florida (Sharp et al., 2020; Neely et al., 2021b) and the U.S. Virgin Islands (Meiling et al., 2020), stony coral tissue loss disease incidence decreases substantially during bleaching events. The relationship between thermal bleaching and disease thus appears to be dependent on the disease type, and fine-scale temporal monitoring is important in assessing the relationship between the stressors.

In addition to immediate post-bleaching losses to disease, longer-term impacts of bleaching on the coral population are well documented through reduced reproductive output of bleached corals (Szmant and Gassman, 1990; Mendes and Woodley, 2002; Levitan et al., 2014). Though the loss of boulder and brain corals through this event was minimal, the ability of these survivors to contribute to the next generation of corals within the next few years is very unlikely. As bleaching events are predicted to become even more frequent in the immediate future, these losses to reproductive capacity are expected to become chronic, with associated losses of juvenile demographic classes.

Other observations that were not quantified during the monitoring surveys but which represented clear ecosystem changes include the decimation of octocorals at nearshore sites. In addition to the near or total bleaching of hard corals, octocorals at all sites showed significant or total bleaching, as did corallimorphs, anemones, and zoanthids. As octocorals, particularly the seafan *Gorgonia ventilina* and the sea plume *Antillogorgia americana* are incredibly abundant and make up a notable component of the benthos, changes in their populations are obvious even when not quantified (Figure S1). At offshore sites, these octocorals appear to have undergone minimal mortality, with most sites having fully-colored individuals and few if any obvious skeletal remains after bleaching of stony corals subsided. Inshore sites on the other hand experienced near decimation of these populations. At Newfound Harbor, Marker 48, and Cheeca Rocks in particular, anecdotal observations suggest that few if any of these individuals survived. We have continued to opportunistically survey small sections of these reefs looking for survivors or new recruits, but to date have found none. The loss of these octocorals, as well as potentially other more rare benthic organisms, may also have ecosystem impacts. For example, Neely (2023) documented unusual basket star behavior following this octocoral die-off. Though stony corals are well studied for their responses to disturbance events, other benthic biota, and the fauna that depend on them, are not. Pre-2023 data do not exist for most of these organisms, and so how bleaching events may impact their behavior or populations remains unknown.

### Conclusions

1. Despite 100% bleaching rates, bleaching-related mortality of brain and boulder corals across the Florida Keys were minimal during the 2023 MHW.
2. The few sites where bleaching-related mortality on brain and boulder corals did occur also experienced the most extreme cumulative thermal stress. At one site, mortality began at relatively low degree heating weeks, suggesting that corals reached thermal maxima rather than death by cumulative thermal stress.
3. At the few sites that did exhibit substantial mortality, brain corals fared substantially worse than boulder corals, with *Pseudodiploria* species, *D. labyrinthiformis*, and *C. natans* having the highest mortality rates. This may have implications for management decisions on outplanting. Anecdotal losses of essentially all Acroporids at the monitored sites may also warrant consideration in whether outplanting programs focusing on susceptible species are sustainable or a worthwhile use of resources.
4. The health status of corals before bleaching had limited impact on how they fared through the event. For one species (*C. natans*), the presence and treatment of SCTLD lesions in June resulted in significantly greater propensity for bleaching-related mortality and black band disease in August, but for all others it did not.
5. The appearance of unknown lesions on *O. faveolata* at some sites as zooxanthellae returned to corals warrants further study. Though the lesions were short-lived, they did cause substantial mortality on some of the largest colonies, and may be an emerging threat.
6. Fate-tracking large numbers of corals across multiple species and reefs provides high-resolution data on survival, recovery, and how health histories evolve across space and time. To our knowledge, this is the only such program in the Florida Keys, and is likely to become increasingly important in understanding the impact of future disturbances across species and reefs.
7. Despite generally high survivorship of non-branching corals, significant legacy effects for the community are likely. These may include reduced coral reproductive output, increased disease susceptibility, and altered benthic community composition. Future monitoring should expand its scope to be able to capture these greater sub-lethal and community level effects if we are to predict the fate of coral reef communities in the Anthropocene.

## Acknowledgements

Data were collected as part of ongoing monitoring for SCTLD intervention work, funded by Florida Department of Environmental Protection’s Coral Protection and Restoration Program, PO C22220. Additional effort was conducted pro bono by the authors. All monitoring was conducted under Florida Keys National Marine Sanctuary permit FKNMS-2020-077. We are grateful to past members of the Neely lab, whose efforts in tagging and treating corals since 2019 made this work possible.

## Supplementary Material

**Figure S1:**
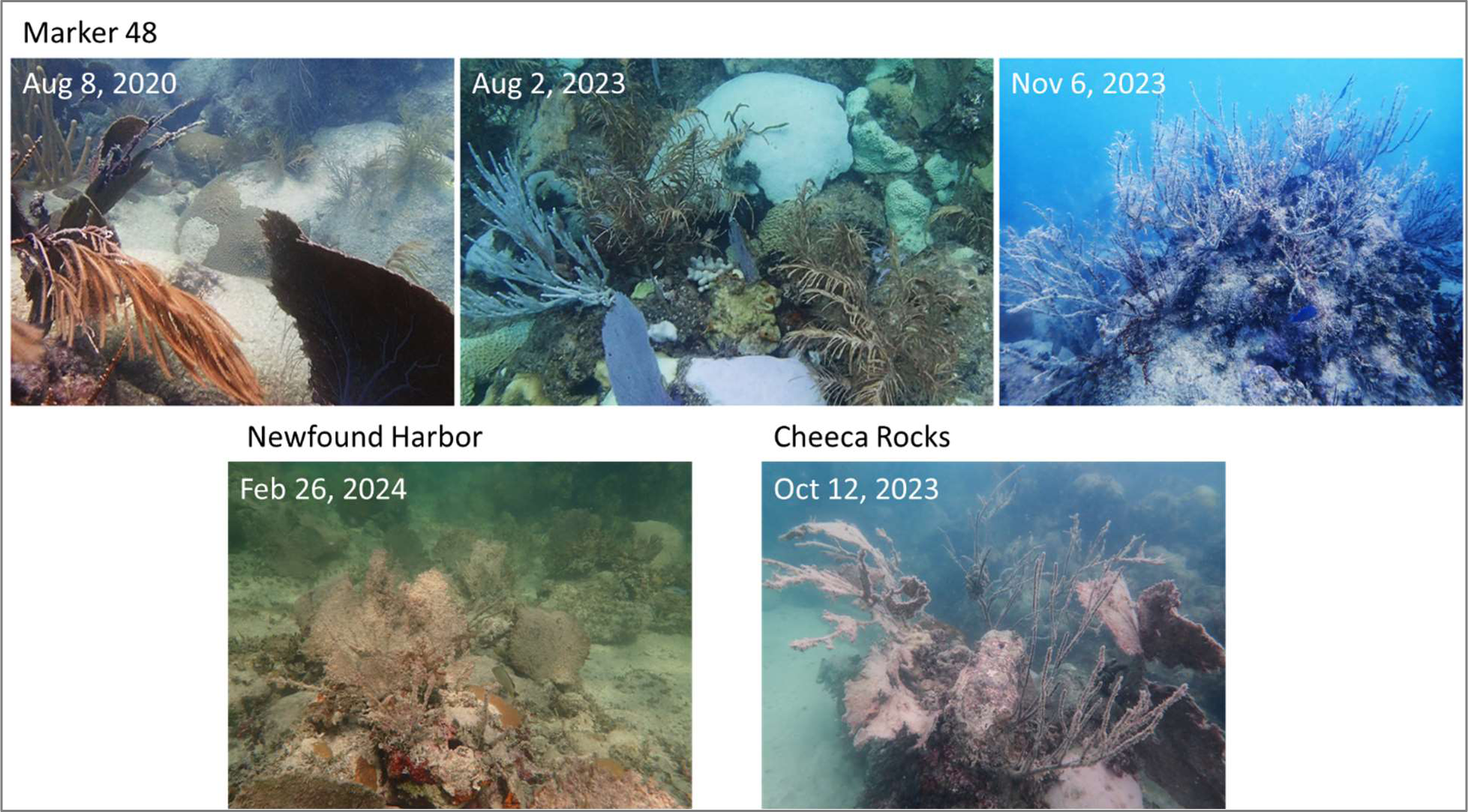
Imagery of inshore reef octocoral dieoffs from the 2023 Florida marine heatwave. Top images are all from the Marker 48 patch reef, showing prevalent live octocorals in 2023, a mixture of unbleached, bleached, and dying octocorals in August 2023, and full mortality of octocorals by November 2023. Bottom images show similarly dead octocoral populations following the MHW at two other inshore patch reefs – Newfound Harbor and Cheeca Rocks.

## References

1. Aragon, T.J., Fay, M.P., Wollschlaeger, D., and Omidpanah, A. (2020). epitools: Epidemiology tools. R package version 0.5*-*10.1.

2. Brandt, M. (2009). The effect of species and colony size on the bleaching response of reef-building corals in the Florida Keys during the 2005 mass bleaching event. Coral Reefs 28, 911–924.

3. Brown, B.E. (1997). Coral bleaching: causes and consequences. Coral Reefs 16, S129–S138.

4. Correa, A., Brandt, M., Smith, T., Thornhill, D.J., and Baker, A. (2009). *Symbiodinium* associations with diseased and healthy scleractinian corals. Coral Reefs 28, 437–448.

5. Fitt, W.K., Spero, H.J., Halas, J., White, M.W., and Porter, J.W. (1993). Recovery of the coral *Montastrea annularis* in the Florida Keys after the 1987 Caribbean “bleaching event”. Coral Reefs 12, 57–64.

6. Gintert, B.E., Manzello, D.P., Enochs, I.C., Kolodziej, G., Carlton, R., Gleason, A.C., and Gracias, N. (2018). Marked annual coral bleaching resilience of an inshore patch reef in the Florida Keys: A nugget of hope, aberrance, or last man standing? Coral Reefs 37, 533–547.

7. Glynn, P.W. (1993). Coral reef bleaching: ecological perspectives. Coral Reefs 12, 1–17.

8. Grajales, A., and Sánchez, J.A. (2016). Holobiont assemblages of dominant coral species (Symbiodinium types and coral species) shape Caribbean reef community structure. Revista de la Academia Colombiana de Ciencias Exactas, Físicas y Naturales 40, 300–311.

9. Hoegh-Guldberg, O. (1999). Climate change, coral bleaching and the future of the world’s coral reefs. Marine and Freshwater Research 50, 839–866.

10. Hothorn, T., Bretz, F., and Westfall, P. (2008). Simultaneous inference in general parametric models. Biometrical Journal 50, 346–363.

11. Kayanne, H. (2017). Validation of degree heating weeks as a coral bleaching index in the northwestern Pacific. Coral Reefs 36, 63–70.

12. Lajeunesse, T.C. (2002). Diversity and community structure of symbiotic dinoflagellates from Caribbean coral reefs. Marine Biology 141, 387–400.

13. Leggat, W.P., Camp, E.F., Suggett, D.J., Heron, S.F., Fordyce, A.J., Gardner, S., Deakin, L., Turner, M., Beeching, L.J., and Kuzhiumparambil, U. (2019). Rapid coral decay is associated with marine heatwave mortality events on reefs. Current Biology 29, 2723–2730. e2724.

14. Levitan, D.R., Boudreau, W., Jara, J., and Knowlton, N. (2014). Long-term reduced spawning in *Orbicella* coral species due to temperature stress. Marine Ecology Progress Series 515, 1–10.

15. Loya, Y., Sakai, K., Yamazato, K., Nakano, Y., Sambali, H., and Van Woesik, R. (2001). Coral bleaching: the winners and the losers. Ecology Letters 4, 122–131.

16. McClanahan, T. (2004). The relationship between bleaching and mortality of common corals. Marine Biology 144, 1239–1245.

17. McWilliams, J.P., Côté, I.M., Gill, J.A., Sutherland, W.J., and Watkinson, A.R. (2005). Accelerating impacts of temperature-induced coral bleaching in the Caribbean. Ecology 86, 2055–2060.

18. Meiling, S., Muller, E.M., Smith, T.B., and Brandt, M.E. (2020). 3D Photogrammetry Reveals Dynamics of Stony Coral Tissue Loss Disease (SCTLD) Lesion Progression Across a Thermal Stress Event. Frontiers in Marine Science 7:597643.

19. Mellin, C., Brown, S., Cantin, N., Klein-Salas, E., Mouillot, D., Heron, S.F., and Fordham, D.A. (2024). Cumulative risk of future bleaching for the world’s coral reefs. Science Advances 10, eadn9660.

20. Mendes, J.M., and Woodley, J.D. (2002). Effect of the 1995-1996 bleaching event on polyp tissue depth, growth, reproduction and skeletal band formation in *Montastraea annularis*. Marine Ecology Progress Series 235, 93–102.

21. Muscatine, L., Mccloskey, L., and Marian, R. (1981). Estimating the daily contribution of carbon from zooxanthellae to coral animal respiration 1. Limnology and Oceanography 26, 601–611.

22. Neely, K.L. (2023). Habitat shift of a basket star during a coral bleaching event. Marine Biodiversity 54, 2.

23. Neely, K.L., Lewis, C.L., Lunz, K.S., and Kabay, L. (2021a). Rapid Population Decline of the Pillar Coral *Dendrogyra cylindrus* Along the Florida Reef Tract. Frontiers in Marine Science 8.

24. Neely, K.L., Macaulay, K.A., Hower, E.K., and Dobler, M.A. (2020). Effectiveness of topical antibiotics in treating corals affected by Stony Coral Tissue Loss Disease. PeerJ 8, e9289.

25. Neely, K.L., Macaulay, K.A., and Lunz, K.S. (2022). Population trajectory and stressors of *Acropora palmata* sites in the Florida Keys. Frontiers in Marine Science 9.

26. Neely, K.L., Shea, C.P., Macaulay, K.A., Hower, E.K., and Dobler, M.A. (2021b). Short– and Long-Term Effectiveness of Coral Disease Treatments. Frontiers in Marine Science 8.

27. NOAA Coral Reef Watch (2023). Florida 5 km single-pixel virtual station time series graphs [Online]. Available: https://coralreefwatch.noaa.gov/product/vs_single_pixel_exp/florida_keys.php#CheecaRocks [Accessed 2024-07-01].

28. NOAA National Weather Service (2023). “Climatological records for Marathon, FL”. Available: www.weather.gov/media/key/Climate/Records/Daily_Records-Marathon.pdf

29. R Core Team (2022). “R: A language and environment for statistical computing”. (Vienna, Austria: R Foundation for Statistical computing).

30. Sharp, W.C., Shea, C.P., Maxwell, K.E., Muller, E.M., and Hunt, J.H. (2020). Evaluating the small-scale epidemiology of the stony-coral-tissue-loss-disease in the middle Florida Keys. PLOS ONE 15, e0241871.

31. Shilling, E.N., Combs, I.R., and Voss, J.D. (2021). Assessing the effectiveness of two intervention methods for stony coral tissue loss disease on *Montastraea cavernosa*. Scientific Reports 11, 1–11.

32. Smith, K.E., Burrows, M.T., Hobday, A.J., King, N.G., Moore, P.J., Sen Gupta, A., Thomsen, M.S., Wernberg, T., and Smale, D.A. (2023). Biological impacts of marine heatwaves. Annual Review of Marine Science 15, 119–145.

33. Smith, T., Brandt, M., Calnan, J., Nemeth, R., Blondeau, J., Kadison, E., Taylor, M., and Rothenberger, P. (2013). Convergent mortality responses of Caribbean coral species to seawater warming. Ecosphere 4, 1–40.

34. Somerfield, P., Jaap, W., Clarke, K., Callahan, M., Hackett, K., Porter, J., Lybolt, M., Tsokos, C., and Yanev, G. (2008). Changes in coral reef communities among the Florida Keys, 1996–2003. Coral Reefs 27, 951–965.

35. Spadafore, R.E., Fura, R., Precht, W.F., and Vollmer, S.V. (2021). Multi-Variate Analyses of Coral Mortality from the 2014-2015 Stony Coral Tissue Loss Disease Outbreak off Miami-Dade County, Florida. Frontiers in Marine Science, 1299.

36. Sully, S., and Van Woesik, R. (2020). Turbid reefs moderate coral bleaching under climate-related temperature stress. Global Change Biology 26, 1367–1373.

37. Szmant, A.M., and Gassman, N.J. (1990). The Effects of Prolonged Bleaching on the Tissue Biomass and Reproduction of the Reef Coral *Montastrea annularis*. Coral Reefs 8, 217–224.

38. Walker, B.K., Noren, H., Buckley, S., and Pitts, K. (2021). Optimizing stony coral tissue loss disease (SCTLD) intervention treatments on *Montastraea cavernosa* in an endemic zone. Frontiers in Marine Science 8, 746.

39. Williams, D., Miller, M., Bright, A., Pausch, R., and Valdivia, A. (2017). Thermal stress exposure, bleaching response, and mortality in the threatened coral *Acropora palmata*. Marine Pollution Bulletin 124, 189–197.

40. Williams, D.E. (2024). “The status of Acropora palmata founders in south Florida after the 2023 marine heatwave”. (NOAA Fisheries).

